# Lung microvascular rarefaction impairs pulmonary gas exchange and exacerbates heart failure with preserved ejection fraction

**DOI:** 10.64898/2026.03.05.709974

**Authors:** Ceren Koçana, Lara Jaeschke, Alexandra Maria Chitroceanu, Qi Zhang, Niklas Hegemann, Pengchao Sang, Qiuhua Li, Mariya M. Kucherenko, Kristin Kräker, Kristina Franz, Alexandr Melnikov, David Faidel, Leonardo A. von der Ohe, Paul-Lennart Perret, Jonathan L. Gillan, Annika Winkler, Edwardo Reynolds, Andreas Kind, Lucie Kretzler, Daniela Zurkan, Veronika Zach, Samuel Mathew Tharakan, Mohammad Amiruddin Hashmi, Bakhrom K. Berdiev, Saba Al Heialy, Mohammed Uddin, Christoph Knosalla, Frank Edelmann, Ralf Dechend, Gabriele G. Schiattarella, Szandor Simmons, Christina Brandenberger, Jana Grune, Wolfgang M. Kuebler

## Abstract

**Background:** Dyspnea and exercise intolerance are the primary clinical symptoms of heart failure. Heart failure patients experience frequent hypoxemic episodes, yet underlying mechanisms and relevance remain poorly understood. In a cohort of heart failure patients and multiple animal models, we identify pulmonary capillary rarefaction driven by excessive autophagy in endothelial cells as a novel mechanism of hypoxemia and cardiac disease progression.

**Methods:** A cohort of heart failure with preserved ejection fraction (HFpEF) patients was analyzed for parameters of left ventricular (LV) dysfunction and pulmonary gas exchange. Morphological and cellular mechanisms of impaired pulmonary oxygenation were assessed in three animal models of heart failure, namely two HFpEF models, SU5416-treated ZSF1 obese rats and high fat diet/L-NAME treated mice, and in rats subjected to aortic banding. Lung microvascular rarefaction was quantified by micro-computed tomography, stereology, flow cytometry and dye efflux. Cellular mechanisms of capillary loss were analyzed by single-cell transcriptomics, electron microscopy and immunofluorescence, and in mice with endothelial-specific deletion of the autophagy gene *Atg7 (Atg7*^EN-KO^).

**Results:** In 234 HFpEF patients, advancing NYHA class was associated with progressive worsening of arterial oxygen saturation at rest and during exercise and a reduced lung diffusing capacity. Impaired gas diffusion correlated with indices of LV diastolic dysfunction. Impaired oxygenation and reduced exercise capacity were similarly evident in animal models of left heart disease, which showed a distinct loss of pulmonary microvessels and capillaries. Lung microvascular endothelial cells in HFpEF showed characteristics of increased autophagic flux and apoptosis. Relative to their wild type HFpEF controls, *Atg7*^EN-KO^ mice had less capillary loss, restored normoxemia, improved exercise tolerance, and mitigated LV diastolic dysfunction. Additional studies in HFpEF mice corroborated the functional relevance of impaired gas exchange for the progression of left heart disease by demonstrating that additional hypoxia aggravated, whereas moderate hyperoxia improved LV function.

**Conclusion:** Our findings identify pulmonary microvascular rarefaction as a novel pathomechanism in heart failure that i) contributes to dyspnea and exercise intolerance, ii) impairs pulmonary gas exchange and iii) accelerates LV disease progression. Strategies targeting this axis such as moderate oxygen therapy may mitigate cardiopulmonary morbidity in heart failure.

**Clinical Trial Registration:** Registered in the DRKS (Deutsches Register für klinische Studien) as trial# DRKS00032974 at https://drks.de/search/en/trial/DRKS00032974.

## Introduction

Heart failure with preserved ejection fraction (HFpEF), defined by a preserved left ventricular (LV) ejection fraction (LVEF) ≥ 50%, accounts for >50% of all heart failure cases globally^1^. It is characterized by diastolic LV dysfunction, increased LV myocardial stiffness and abnormal ventricular-vascular coupling. HFpEF is commonly associated with and driven by characteristic comorbidities including hypertension, metabolic syndrome (obesity, diabetes mellitus, dyslipidemia), chronic kidney disease as well as aging^2^. In spite of the recent introduction of SGLT2 inhibitors as the first approved clinical therapy for HFpEF (EMPEROR-preserved trial, DELIVER trial), morbidity and mortality remain high, stressing a major unmet clinical need^3,4^.

Besides fatigue and impaired exercise capacity, dyspnea on exertion or even at rest is the cardinal clinical symptom in HFpEF patients. Limitations in exercise capacity can be evidently attributed to a reduced ability to increase cardiac output in HFpEF patients, which is often normal at rest^5^. Yet, dyspnea is not a direct consequence of a reduced cardiac reserve but rather a sign of impaired arterial blood oxygenation and/or impaired lung inflation as measured by arterial chemosensors and pulmonary stretch sensors, respectively. Indeed, several clinical studies show evidence of systemic hypoxemia in HFpEF patients. Specifically, oxygen desaturations with SaO_2_ values <90% are 3 times more frequent and last 5-6 times longer in HFpEF patients as compared to healthy controls^6^. In a study of 539 HFpEF patients undergoing invasive cardiopulmonary exercise testing, exertional hypoxemia with SaO_2_ <94% was observed in 25% of patients, and particularly prominent in older and more obese patients^7^. Further regression analyses revealed a direct relationship between increased pulmonary capillary pressure and lower arterial oxygen tension, especially during exercise^7^. Importantly, impaired arterial oxygenation cannot be attributed to LV-failure associated hemodynamic changes *per se*, as reduced cardiac output and the resulting lower blood flow velocity across the pulmonary microvasculature will improve, rather than impair, oxygen uptake, while pulmonary congestion increases vascular surface area and thereby facilitates enhanced diffusion. Rather, systemic hypoxemia likely reflects impaired oxygen uptake in the lung. Consistent with this view, clinical studies show a marked increase in the alveolar-arterial oxygen difference in patients with HFpEF^7^ or acute heart failure^8^, respectively. Similarly, the lunǵs diffusing capacity for carbon monoxide (DLCO) is characteristically reduced in HFpEF patients^9,10^ suggesting impaired alveolo-capillary gas exchange. Clinical HFpEF is frequently associated with extensive vascular remodeling in both pulmonary arteries and veins^11,12^, yet, HFpEF-associated structural changes at the level of the alveolo-capillary interface as the primary functional unit of pulmonary gas exchange remain largely unclear. In the present study, we hypothesized that HFpEF is not only associated with pulmonary arterial and venous remodeling, but also with extensive structural remodeling of pulmonary microvessels resulting in reduced alveolo-capillary gas exchange. The ensuing hypoxemia will in turn aggravate right ventricular (RV) and LV dysfunction, thereby propagating HFpEF deterioration in a positive feedback loop.

## Methods

### Patient data

234 patients with HFpEF were prospectively recruited into the clinical cohort of the Collaborative Research Center 1470 (CRC1470), a single-center consortium supported by the German Research Foundation (DFG, Deutsche Forschungsgemeinschaft). The study complied with the Declaration of Helsinki and was approved by the Ethics Committee of Charité – Universitätsmedizin Berlin (Berlin, Germany; approval number EA1/224/21). All participants were older than 18 years and provided written informed consent. A positive diagnosis of HFpEF according to the current American College of Cardiology (ACC) criteria was required for inclusion in the cohort, and defined as follows: (i) symptoms and/or signs of heart failure corresponding to at least New York Heart Association (NYHA) class II prior to medication; (ii) a left ventricular ejection fraction (LVEF) exceeding 50%; (iii) objective evidence of cardiac structural and/or functional abnormalities indicative of diastolic dysfunction or increased left ventricular filling pressures. Structural abnormalities were defined as a left ventricular mass index (LVMI) > 95 g/m² in females or > 115 g/m² in males, a relative wall thickness (RWT) > 0.42, or a left atrial volume index (LAVI) > 34 mL/m². Functional abnormalities included an E/e′ ratio at rest > 13 or a pulmonary artery systolic pressure (PASP) > 35 mmHg. Inclusion further required plasma levels of N-terminal pro–B-type natriuretic peptide (NT-proBNP) > 125 pg/mL. Exclusion criteria included a life expectancy of less than one year, a history of heart failure with mid-range or reduced ejection fraction, acute coronary syndrome within the preceding 30 days, cardiac surgery within the past three months, or end-stage renal disease with or without hemodialysis. Further exclusions comprised severe valvular heart disease, hypertrophic cardiomyopathy, amyloidosis, congenital heart disease, sarcoidosis, and constrictive pericarditis. Patients with extracardiac disorders that could account for their symptoms, such as chronic obstructive pulmonary disease (COPD) GOLD stage > 2, pulmonary arterial hypertension, or moderate to severe anemia (hemoglobin < 10 g/dL in males and < 9.5 g/dL in females), were also excluded. All participants underwent a comprehensive clinical assessment, including two-dimensional echocardiography (2DE) of both ventricles, spiroergometry, diffusion single breath analysis, and standard blood work.

### Animals

All animal experiments were performed in compliance with the Directive 2010/63/EU of the European Parliament, the German Animal Welfare Act, and the Guide for the Care and Use of Laboratory Animals (NIH Publication No. 85-23, revised 1985). Experimental procedures were approved by the local regulatory authority (Landesamt für Gesundheit und Soziales, Berlin, Germany; protocol numbers G0008/22, G0025/24, G0030/18 and G0085/23) and adhered to the Animal Research: Reporting of In Vivo Experiments (ARRIVE) guidelines.

All animals were housed for at least 1 week to acclimatize to laboratory conditions before starting experimental procedures. Two to four animals were housed per cage, divided by sex and treatment. Animals were kept in a 12-h:12-h light/dark cycle with free access to food and water in individually ventilated cages and pathogen-free rooms. All animals underwent weekly body weight and daily health monitoring.

*C57BL/6J* mice and Sprague-Dawley rats were purchased from Janvier-Labs. ZSF1 rats were purchased from Charles River Laboratories. Mice with an endothelial cell (EC)-specific deletion of the essential autophagy gene *Atg7* were generated by crossing *Atg7*^flox/-^ mice with mice constitutively expressing Cre recombinase under control of the VE-cadherin promoter (*Cdh5*-Cre^tg/+^). *Atg7*^flox/flox^; *Cdh5*-Cre^tg/+^ mice were used as EC-specific *Atg7* knockout mice (*Atg7*^EN-KO^), with *Atg7*^flox/flox^; *Cdh5*-Cre^+/+^ (*Atg7*^EN-WT^) mice serving as corresponding controls. Ear biopsies were collected for genotyping by PCR. Animals were randomly assigned to either control, heart failure, or intervention groups specified below.

*Murine HFpEF model.* Eight week-old *C57BL/6J* mice of both sexes were divided into four experimental groups: i) naive mice receiving standard diet (chow, 9% fat, Ssniff Spezialdiäten GmbH, Soest, Germany) and regular drinking water serving as controls, ii) mice receiving a rodent diet with 60% fat (high-fat diet – HFD, Ssniff Spezialdiäten GmbH, Soest, Germany) and N[ω]-nitro-l-arginine methyl ester (L-NAME, 0.5 g/L, N5751-10G, Sigma-Aldrich) for twelve weeks to induce HFpEF^13^, iii) mice receiving HFD, L-NAME diluted in drinking water and additional exposure to chronic hypoxia (10% O_2_) for two weeks from week 10 to assess the effects of hypoxemia on LV function, iv) mice receiving HFD, L-NAME diluted in drinking water and additional exposure to moderate hyperoxia (40% O_2_) for two weeks from week 10 to restore normoxemia in HFpEF mice and assess effects on RV and LV function. All groups were age– and sex-matched and run in parallel.

*Rat HFpEF model.* To induce HFpEF in rats, male obese ZSF1 rats (8– to 10-weeks of age) received a single 100 mg/kg subcutaneous injection of the vascular endothelial growth factor receptor-2 antagonist Sugen (Semaxinib, SU5416, MedChemExpress, HY-10374, Monmouth Junction, NJ) dissolved in CMC buffer (0.5% sodium carboxymethyl cellulose, 0.4% polysorbate 80, 0.9% sodium chloride, and 0.9% benzyl alcohol) as described^14^. Lean ZSF-1 rats served as controls. Endpoint analyses were performed after 14 weeks.

*Rat heart failure model.* As additional model, we induced left heart failure in 4– to 8-weeks old male Sprague–Dawley (SD) rats (∼100 g body weight) by aortic banding (AoB) surgery under ketamine/xylazine (87/13 mg/kg body weight) anesthesia as previously described^15,16^. In brief, a metal clip with an open inner diameter of 0.8 mm was surgically implanted across the aorta just after the coronary arteries branch off. Sham animals were subjected to the same surgical procedure without clip placement. Endpoint examinations were performed at weeks 3 and 9 after surgery. Researchers performing endpoint and post-mortem analyses were blinded for groups and treatment.

### Treadmill exercise

Mice were initially acclimatized to a treadmill (CaloTreadmill, TSE Systems, Germany) by running uphill (20°) at a constant speed (5 m/min) for 5 min for 3 consecutive days. After acclimatization, an exhaustion test was performed by running uphill (20°) on the treadmill at warm-up speed (5 m/min) for 4 min and then increasing the speed to 14 m/min for 2 min. Every 2 min thereafter, the speed was increased by 2 m/min until the animal was exhausted. Exhaustion was defined by two separate criteria: (i) the animal was exposed to the shock grid at the back of the treadmill for ≥10 seconds or (ii) the animal spent less than 2 s on the treadmill before contacting the shock grid again (i.e. lack of motivation). Running distance and the maximum rate of oxygen consumption (V̇O_2_ max) were assessed using PhenoMaster software V7.1.6 (TSE Systems, Germany). Arterial oxygen saturation (SaO₂) was measured non-invasively immediately before and after the exercise protocol using a MouseOx Plus collar sensor system (STARR Life Sciences, Oakmont, PA, USA).

### Echocardiography

Progressive cardiac dysfunction was monitored by non-invasive small animal echocardiography according to the standards defined by the American Society of Echocardiography (ASE). To this end, animals were anesthetized with 1-2% (mice) / 1.5-3% (rats) isoflurane inhalation anesthesia. Eye ointment was applied to protect the animalś eyes from drying. Echocardiographic imaging was performed using ultrahigh frequency linear array transducers (MX400, 20–46 MHz for mice; MX250, 15–30 MHz for rats) on a Vevo^®^ 3100 high-resolution imaging system (FUJIFILM VisualSonics Inc., Toronto, ONT, Canada). In brief, animals were fixed in supine position on a heated table and the thorax was depilated (Veet depilatory cream and – for rats – shaving). The heart was visualized parasternally in the short axis, the long axis and in apical four-chamber view. In addition, measurements were taken in brightness (B)– and motion (M)-Mode (short axis and long axis) to calculate the LV and RV chamber dimensions at the level of the papillary muscles. Doppler imaging was recorded for the LV and the pulmonary artery, and speckle tracking echocardiography was performed for LV strain analyses. All images were stored as raw data in DICOM format for offline analysis.

Image analysis was performed using the VevoLAB Version 5.7.1 (FUJIFILM VisualSonics). Ejection fraction, stroke volume, end-diastolic volume, and heart rate were determined from B-mode images of the LV in the parasternal long-axis view using the LV Trace tool. End-diastolic diameter was obtained from M-mode images in the same view with the semi-automatic LV Trace function. Two-dimensional speckle-tracking echocardiography for assessing global longitudinal peak strain (GLS) was carried out using VevoStrain 2.0 software (FUJIFILM VisualSonics) and a speckle-tracking algorithm. Semi-automated endocardial border tracing was carried out on three consecutive cardiac cycles per animal, selected from a B-mode cine loop obtained in the parasternal long-axis view at end-systole. B-mode images were chosen based on frame rates above 200 fps and clear visualization of endocardial and epicardial LV borders. Peak GLS values were derived from six independent anatomical LV segments.

### Invasive hemodynamics

Mice were anesthetized by intraperitoneal (i.p.) injection of ketamine (120 mg/kg body weight) and xylazine (16 mg/kg body weight), and neck and chest were depilated as described above. Mice were then placed on a specially prepared, sterile operating table equipped with a heating mat. After checking the interdigital reflex to ensure a sufficient depth of anesthesia, the jugular vein and the carotid artery were sequentially catheterized with a 1F microtip Millar catheter and the catheter was advanced into the RV and LV for measurement of RV systolic pressure (RVSP) and LV systolic pressure (LVSP), respectively.

Rats were anaesthetized by i.p. ketamine (87 mg/kg body weight) and xylazine (13 mg/kg body weight), neck and chest were depilated as described above, and animals were placed on a sterile operating table as reported for mice. After ensuring sufficient depth of anesthesia by checking the inter-phalangeal reflex, rats were tracheotomized and mechanically ventilated as follows: The trachea was visualized with a cervical midline incision and cannulated. Lungs were perioperatively ventilated at a breathing rate of 90 breaths/min and a tidal volume of 8.5 mL/kg body weight. Following thoracotomy by vertical sternotomy, cardiac catheterization was performed directly through the apex of the heart, and LVSP and RVSP were recorded via a Millar catheter positioned in the left and right ventricle, respectively. Stable pressure profiles were recorded for at least 2 min by PowerLab 4/35 connected to LabChart v. 8 software (ADInstruments, Sydney, Australia) and analyzed off-line by an independent observer blinded to the respective protocol. LVSP and RVSP were calculated as the mean systolic pressure measured over 10 s in hemodynamically stable recordings. Arterial and venous blood samples were obtained for blood gas analyses. Animals were then euthanized under deep anesthesia by exsanguination, and blood and tissue samples were collected.

### Gravimetry

Following euthanasia, the heart was excised after perfusion through the left ventricle with 10 mL ice-cold 1X PBS, the ventricles were separated from the atria and dissected into RV and LV including septum (LV+S). RV and (LV+S) weight were determined by gravimetry and normalized to body weight (for rats) and tibia length (for mice), respectively.

### Flow cytometry

Lung tissue was subjected to enzymatic digestion using collagenase I (450 U/mL), collagenase XI (60 U/mL), DNAse I and hyaluronidase (60 U/mL) (Sigma-Aldrich) buffered in HEPES buffer (Corning) and 1x PBS for 30 min at 37°C and 700 rpm. Cells were filtered through a 40-µm nylon mesh for mouse lungs and a 70-µm nylon mesh for rat lungs, washed, and centrifuged (5 min, 350xg, 4°C) to obtain single-cell suspensions. Red blood cells were lysed with 1x red blood cell lysis buffer (BioLegend).

Cell suspensions from mouse lungs were stained at 4°C for 30 min with CD45-Brilliant Violet 711 (clone 30-F11, 1:300, 103147, BioLegend), CD31-Alexa Fluor 700 (clone 390, 1:600, 102443, BioLegend) antibodies and Alexa Fluor 647-conjugated isolectin griffonia simplicifolia IB4 (GS-IB4, 1:1000, I32450, Invitrogen) with Fc block by CD16/CD32 antibody (clone 2.4G2, 1:600, BD Bioscience, 553141). Viable cells were detected as negative for DAPI staining (BD Bioscience, 564907).

Cell suspensions from rat lungs were stained at 4°C for 30 min with CD45-PerCP/Cyanine5.5 (clone OX-33, 1:800, 202318, BioLegend), CD31-PE (clone TLD-3A12, 1:800, 555027, BD Pharmingen) antibodies and Alexa Fluor 647-conjugated isolectin griffonia simplicifolia IB4 (GS-IB4, 1:1000, I32450, Invitrogen) with Fc block by CD32 antibody (clone D34-485, 1:1000, BD Pharmingen, 550271). Viable cells were detected as negative for 7-AAD viability staining (BioLegend, 420404).

Endothelial cells were identified as CD45^-^CD31^+^ cells, and lung microvascular endothelial cells were subsequently gated as CD31^+^GS-IB4^+^ cells. Autophagic flux was assessed with an Autophagy detection kit (Abcam, ab139484) using monodansylcadaverine as fluorescent marker for autophagic vacuoles. All flow cytometric data were recorded on a Symphony A5 flow cytometer with FACSDiva 8.0.1 software and analyzed with FlowJo 10.8.1 software (BD Biosciences).

### Stereology

Post-mortem, murine and rat lungs were perfused with fixative (1.5% glutaraldehyde and 1.5% paraformaldehyde (PFA) in 0.15 Mol/L HEPES buffer; pH 7.35) at an initial inflation pressure of 25 cmH_2_O and then at 20 cmH_2_O during fixation. Lungs and heart were removed and transferred to the same fixative solution and then further processed for stereological imaging. After separation of heart and lungs, volumes of the left and right lung [V(lung)] were determined separately by fluid displacement according to Archimedes’ principle^17^. The lung tissue was embedded into 4% agar and then cut into slices with an isotropic uniformly random (IUR) protocol using an orientator^18^. Every second section was embedded in glycol methacrylate (Technovit^®^ 7100, Kulzer GmbH) as described previously^19^. The remaining tissue slices were further cut into 1 mm^3^ tissue blocks and embedded in Epon resin (Roth, Karlsruhe, Germany) as described before^20^. Tissue sections embedded in Technovit^®^ were cut into 1.5 µm thick sections, stained with toluidine blue and imaged on a Axio Scan.Z1 slide scanner (Zeiss, Göttingen, Germany). From these, the volume of the lung parenchyma was assessed using the newCAST stereology software (Visiopharm, Horsholm, Denmark). An objective lens magnification of 20x with a test grid consisting of 3 x 3 points was used to count the number of points hitting parenchyma (P_Par_) and nonparenchymal (P_NonPar_) structures and to calculate the total volume of parenchyma of the lung (V(par,lung)) by multiplying P_Par_ with V(lung).

The total number of capillaries was quantified according to a standard protocol described by Willführ and colleagues^21^ by estimating the Euler number of capillary networks. To this end, the disector principle was applied^21^ and 1 µm thick consecutive sections were cut from Epon embedded tissue blocks stained with toluidine blue for parenchyma and nonparenchymal structures. Sections were again digitalized with the AxioScan Z.1 slide scanner at 40 x magnification. The Euler-Poincaré characteristic/Euler number of capillary networks was calculated by counting the topological constellations: new, isolated parts (islands); enclosed cavities within an existing profile (holes); and in the majority of cases in this study, extra connections between two isolated structures (bridges). The first and third sections were analyzed using the disector method^22^, and stereological estimation was performed by Visiopharm software. In brief, tissue sections were digitally aligned, and profile events were estimated within an unbiased counting frame with an area of 9123.42 μm2 (A(frame)). The reference volume for the profile estimation V(dis) was calculated from the number of counting frames (P(ref)) that were assessed and the height of the disector pairs (h, 2 µm) as shown below

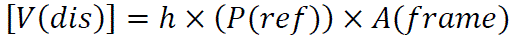

From the raw counts, the Euler number of capillaries [χ(cap)] was determined as:

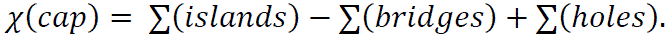

The numerical density [Nv(cap/lung)] was then calculated using χ(cap) as well as the total number of capillaries per lung [N_V_(cap,lung)]:

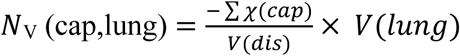

In a subset of experiments (specifically, in the SU5416-treated ZSF1 obese rats and their corresponding controls), image quality and the resolution at light microscopic level was not sufficient to allow for the stereological approach outlined above. Instead, we utilized TEM images from these lungs to calculate total capillary volume density within the inter-alveolar septa of the lung as previously described^20^. In brief, a point grid test system was superimposed onto the digitized TEM images to estimate volume densities of the respective compartments according to design-based stereological principles. By point counting, the epithelial, capillary endothelial, interstitial and capillary lumen (P_capillary lumen_) compartment were quantified using STEPanizer software^23^.The total number of test points hitting the alveolar septum (P_total_) was defined as the sum of points hitting alveolar epithelium, capillary endothelium, interstitium, and capillary lumen. Total capillary volume density within the septum was calculated as: Vv(cap/sept) = P_capillary lumen_ / P_total_.

### Transmission electron microscopy

For transmission electron microscopic (TEM) analysis, ultrathin sections of 70 nm were cut from Epon embedded tissue blocks collected on pioloform-coated copper grids and stained with uranyl acetate and lead citrate60. TEM images were acquired by a Zeiss EM 906 electron microscope at 80 kV acceleration voltage (Carl Zeiss, Oberkochen, Germany) equipped with a 2K CCD camera (TRS, Moorenweis, Germany). To assess autophagic stress in EC, autophagosomes were identified as double membrane structures containing undegraded cytoplasmic components such as endoplasmic reticulum membranes, mitochondria, or ribosomes as previously described^24^.

### Micro-computed tomography

To prepare lung tissue for micro-computed tomographic (µCT) imaging of the microvascular bed, the contrast agent Vascupaint™ (SKU: MDL-121, MediLumine, Canada) was infused immediately post-mortem into the RV, from where it distributed through the pulmonary vascular system. Once the contrast agent emerged from the LV, perfusion was maintained for an additional 15 min to allow for optimal distribution of the contrast agent across the pulmonary vasculature. Then, the pulmonary artery and the opening in the left ventricle were ligated and the entire body of the animal was placed in the refrigerator (4°C) overnight to harden the Vascupaint inside the lung. The next day, the lungs were transfered to 4% PFA for storage and subsequent scanning by µCT.

Lungs were imaged on a SkyScan 1276 µCT imager (Bruker, Belgium) using the vendor’s software for image acquisition (v.1.4) with the following parameters: X-ray source voltage of 100 kV and source current of 80 µA, step and shoot mode, 360° acquisition, Cu 0.25 mm filter, rotation step of 0.3°, pixel size of 8 µm, exposure time of 2,000 ms, and frame averaging of 4. Flat field correction was always applied.

Images were reconstructed to 8-bit files using NRecon (v.1.7.4.6, Bruker, Belgium). Beam-hardening correction of 12% and ring-artefact correction of 2 were applied. Image analysis was performed with CTAn (v.1.20.8.0, Bruker, Belgium) starting with a delineation of a volume of interest (VOI) to separate the tissue from the surrounding elements in the image. Next, multiple analysis steps were applied to calculate the total organ volume, the volume of the vascular tree and the distributions of the diameters of the segmented blood vessels as follows: Gaussian filtering 3D (radius = 2), global thresholding (11-255), morphological opening 3D (r=3), sweeping all except the largest object (3D), morphological closing 3D (r=3), removing pores 3D, morphological closing 2D (r=8), removing pores 3D, morphological opening 2D (r=8), removing speckles 3D (<5,000 voxels), saving bitmap, 3D analysis, reloading image, median filtering 3D (r=1), global thresholding (62-255), removing speckles 3D (<30 voxels), saving bitmap, and finally 3D analysis with computation of the trabecular thickness distribution. In a few cases the global threshold for segmentation of the total organ volume was set at 15 instead of 11 to achieve the same accuracy. Total tissue volume and total vascular volume were calculated. To specifically capture changes in the pulmonary microvasculature, vascular volume of vessels <250 µm in size was quantified; a vessel size of 110 µm was defined as lower cutoff for reliable resolution of single vessel segments.

### Pulmonary capillary surface area

In a subset of AoB experiments, pulmonary capillary surface area was determined by the blue dextran (BD, Sigma-Aldrich, Germany) efflux method^25^. In brief, lungs were isolated from AoB or sham rats, perfused with Krebs-Ringers buffer containing 5% dextran, and then loaded with BD by addition of 0.67 mg/mL BD to the perfusate. Samples were collected from the outflow reservoir at 1 min intervals, then circulatory flow was re-started with BD-free Krebs-Ringers buffer containing 7% bovine serum albumin (BSA) with sample collection from the outflow reservoir in 30 s intervals. The perfusion volume was recorded, and the BD concentration in each sample was spectrophotometrically determined at λ=630 nm. As the pulmonary capillaries will bind BD during dye loading as a function of their total surface area, the subsequent efflux of dextran during BD-free perfusion can be used to calculate how much BD did initially bind to the capillary bed, and thus to estimate the pulmonary capillary surface area.

### Lung immunohistology

Lung tissue sections were fixed in 4% PFA and stored in 1x PBS. After embedding in paraffin, 5 µm thick sections were cut, dried at 65°C and deparaffinized by Xylol (2X 5 min, Roth) and a decreasing alcohol series (2X 100%, 90%, 80%, 70%) finished with distilled water (5 min each step). Samples were boiled in a microwave at 600W for 10 min in Tris-EDTA buffer, then cooled for 20 min. After rinsing sections for 5 min in PBS, samples were blocked by PBTB (0.2% Triton X-100, 5% normal goat serum, 0.2% BSA in PBS) at room temperature for 1 h and finally incubated with primary antibodies at 4°C overnight. Primary antibodies used for immunostaining of lung tissue sections and respective dilutions were: PECAM-1 (1:400; goat polyclonal; #AF3628, R&D Systems, Minneapolis, MN), cleaved-caspase 3 (Cl-Cas3; 1:400; rabbit polyclonal; #9661, Cell Signaling, Danvers, MA), and cleaved PARP-1 (1:400; Abcam, #ab32064). The tissue sections were washed 1x with PBS, followed by incubation with fluorescent dye-conjugated secondary antibodies at room temperature for 1 h and again 1x wash in PBS. Secondary antibodies were: Alexa Fluor^TM^ 594 rabbit anti-goat antibody (1:500; #A11080), and Alexa Fluor^TM^ 647 goat anti-rabbit antibody (1:500; #A21245, all Invitrogen^TM^, Carlsbad, CA). After 1x wash in PBS, tissue sections were stained with DAPI (1:1000; # D9542, Sigma-Aldrich) for 10 min. Following another 1x wash in PBS, tissue sections were mounted with Fluoromount-GTM without DAPI (Invitrogen^TM^, Carlsbad, CA), imaged by confocal microscopy (Nikon Scanning Confocal A1Rsi+, Tokyo, Japan) and analyzed using ImageJ. Representative images were selected from randomly acquired fields using systematic sampling across each section. For each group, the representative field was chosen from an animal with a quantified value closest to the group median. Display settings were applied identically across groups.

### Single cell RNA sequencing

The dataset used in this study was generated in a previous study from lungs of *C57BL/6J* mice treated with L-NAME and HFD (analogous to our murine HFpEF model) for 2 weeks^26^, and is publicly available through the Gene Expression Omnibus under accession number GSE244309. The full code used for the original scRNA-seq analyses is available at https://github.com/kropskilab/myeloid_il1b. We re-analyzed this single-cell RNA sequencing (scRNA-seq) dataset using the Seurat R package (v4) alongside supporting packages dplyr, tidyr, ggplot2, and viridis for data processing and visualization. The Seurat object was first loaded, and samples were annotated into experimental groups (“Control” or “HFpEF”) based on sample identifiers. Gene expression was normalized using Seurat’s NormalizeData function (log-normalization). For targeted analyses of integrated apoptotic and autophagic markers, curated gene sets related to regulated cell death pathways and autophagy were selected based on existing literature. The applied gene sets are given in Suppl. Tables 1A and 1B. Individual cell types were identified as in the original publication^26^ and expression matrices for the selected gene sets were extracted for specific cell types. For each cell, a “net expression” score was calculated as the sum of expression values across all genes in the set. Individual samples were grouped by experimental condition. Data are displayed as boxplots and scatterplots of net expression.

### Data analyses and statistics

Statistical analyses were performed using GraphPad Prism 9 software (GraphPad Software, Inc., Boston, MA). Results are reported as median with interquartile range (IQR) or mean±SEM. Normality was assessed by Shapiro-Wilk normality test. For 2-group comparisons, parametric data sets were analyzed by unpaired parametric two-tailed t-test; non-parametric data sets by Mann-Whitney U-test. P-values were adjusted for multiple testing using the false discovery rate (FDR) method. For comparisons between multiple groups, one-way analysis of variance (ANOVA) was used for parametric data sets, followed by Dunńs test. For correlations analysis, the Pearson correlation coefficient r was calculated. Statistical significance was assumed at p < 0.05.

## Results

### Impaired oxygenation in HFpEF patients

To assess parameters of pulmonary oxygenation in clinical HFpEF, we first analyzed data from 234 HFpEF patients enrolled in the clinical cohort of the CRC1470 at the German Heart Center of the Charité, Berlin. Patients were stratified based on NYHA classification, and echocardiographic evaluation confirmed increased HFpEF severity – namely a higher ratio of early to late LV filling velocity (E/A), a higher ratio of early diastolic mitral inflow velocity over early diastolic mitral annular tissue velocity (E/é), and a lower global longitudinal strain (GLS) – with higher NYHA class (Fig. 1A). LV ejection fraction (LVEF) did not differ between classes and was consistently >50% confirming the diagnosis of HFpEF. LV end-diastolic volume (EDV) showed a trend to lower values with higher NYHA class that, however, did not reach the level of significance. Complete patient demographics (Suppl. Table 2) and the full set of echocardiographic parameters (Suppl. Table 3) are provided as Supplementary Material.

**Figure 1.**
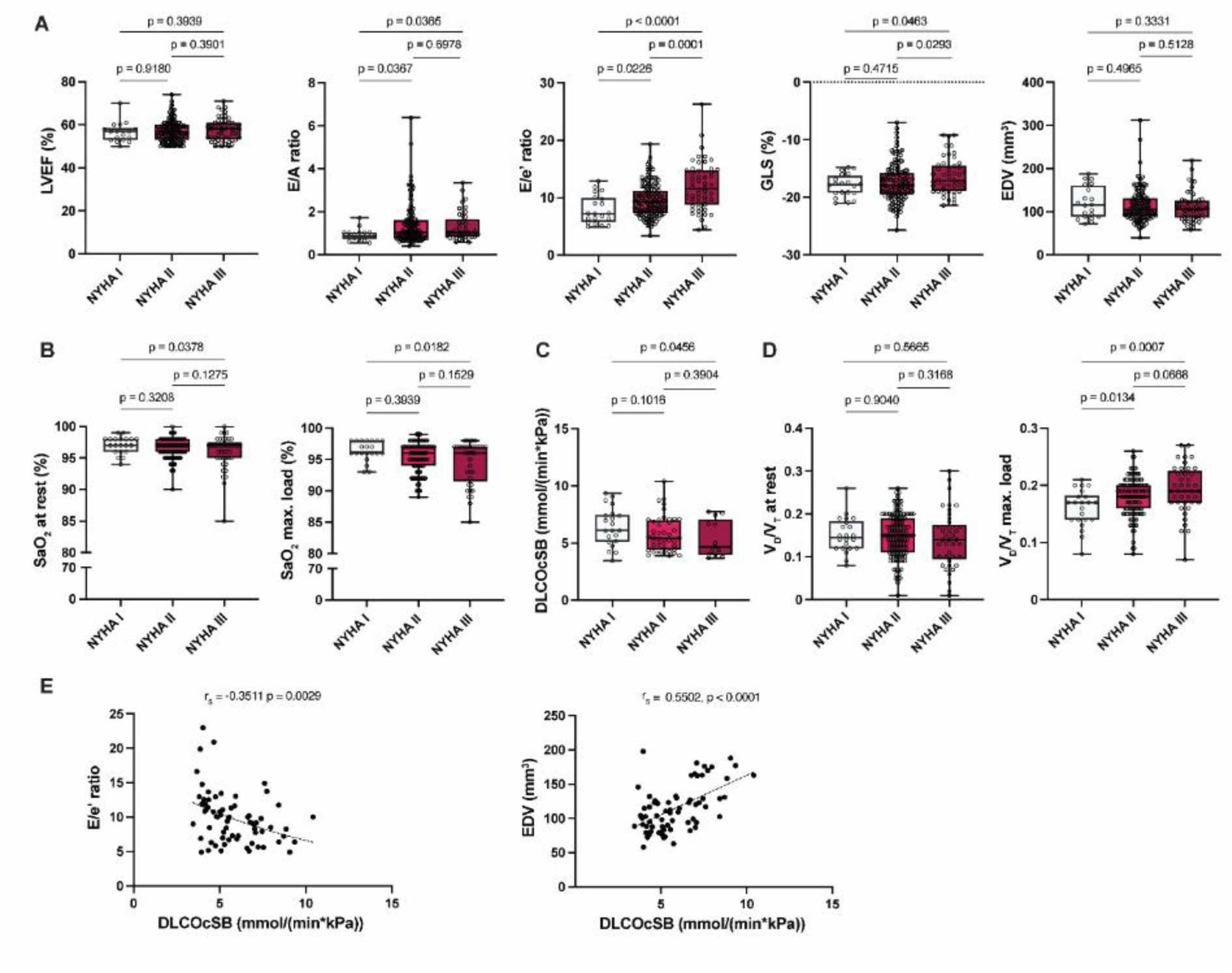
Impaired oxygenation in HFpEF patients. A total of 234 HFpEF patients of the clinical cohort of the CRC1470 were stratified according to New York Heart Association (NYHA) class I (n=25), II (n=157), and III (n=52). **A.** Echocardiographic analysis of left ventricular (LV) ejection fraction (LVEF), ratio of early to late LV filling velocity (E/A ratio), ratio of early diastolic mitral inflow velocity over early diastolic mitral annular tissue velocity (E/e’ ratio), global longitudinal strain (GLS) and end-diastolic volume (EDV); **B.** Arterial oxygen saturation (SaO_2_) at rest and during cardiopulmonary exercise testing at maximal load; **C.** Diffusing capacity of the lungs for carbon monoxide after correction for hemoglobin (DLCOc-SB); **D.** Estimated dead space relative to tidal volume (V_D_/V_T_) at rest and during cardiopulmonary exercise testing at maximal load. **E.** Correlation analysis of E/é ratio and EDV against DLCOc-SB. Dots represent individual patients. Data are presented as median with interquartile range (IQR). Kruskal-Wallis test followed by uncorrected Dunn’s multiple comparisons test; r_s_: Spearman coefficient of correlation; P-values as indicated.

Inversely to HFpEF severity, arterial oxygen saturation (SaO_2_) measured either at rest or during cardiopulmonary exercise testing at maximal load decreased with advancing NYHA class, with patients of NYHA class III showing significantly lower SaO_2_ levels than those with NYHA class I (Fig. 1B). These findings not only consolidate previous reports of systemic hypoxemia in HFpEF patients, but demonstrate a direct association between disease severity and the degree of hypoxemia both at rest and – even more pronounced – during exercise.

Consistent with impaired blood oxygenation during pulmonary transit, assessment of single-breath carbon monoxide uptake in the same patient cohort revealed a decreased diffusing capacity of the lungs (DLCO-SB) in NYHA class III relative to NYHA class I patients (Suppl. Fig. 1A). Alveolar volume (V_A_) and hemoglobin (Hb) concentration, which may affect DLCO, did not differ between groups (Suppl. Fig. 1B,C), suggesting impaired gas diffusion across the alveolo-capillary membrane in severe HFpEF. This notion was confirmed by lower DLCO-SB after correction for Hb (DLCOc-SB; Fig. 1C) and a lower transfer coefficient for CO (KCOc-SB), which corrects DLCOc-SB for V_A_ (Suppl. Fig. 1D). In line with this notion, estimated dead space (V_D_/V_T_) increased with NYHA class in exercise, although it did not differ at rest (Fig. 1D). Finally, DLCOc-SB correlated with parameters of LV diastolic function, specifically with E/é and EDV indicating that impaired gas diffusion in the lung was directly related to LV diastolic dysfunction in HFpEF (Fig. 1E).

### Lung microvascular rarefaction in HFpEF animals

To probe for microvascular rarefaction as potential mechanism of impaired pulmonary gas exchange, we next characterized microvascular remodeling across two different preclinical models of HFpEF. First, we used the established model of SU5416-treated ZSF1 obese rats (Fig. 2A). Consistent with HFpEF, rats showed characteristics of LV diastolic dysfunction, namely an altered E/A ratio and a reduced GLS, while the increase in E/é ratio did not reach the level of significance (Fig. 2B). LV EF remained unchanged at approximately 60%. Invasive hemodynamic measurements demonstrated an increase in RVSP in line with secondary pulmonary hypertension (PH). µCT imaging (a representative recording of which is shown as Suppl. Video 1) revealed a decrease in pulmonary vascular volume that was particularly prominent in vessels <250 µm in diameter (Fig. 2D; Suppl. Fig. 2A), and a decrease in capillary volume density within the interalveolar septa of the lung as determined by stereological assessment of TEM images (Fig. 2E). A full set of echocardiographic, hemodynamic and gravimetric data on SU5416-treated ZSF1 obese rats is given in Suppl. Table 4, and additional stereological parameters are given in Suppl. Table 5.

**Figure 2.**
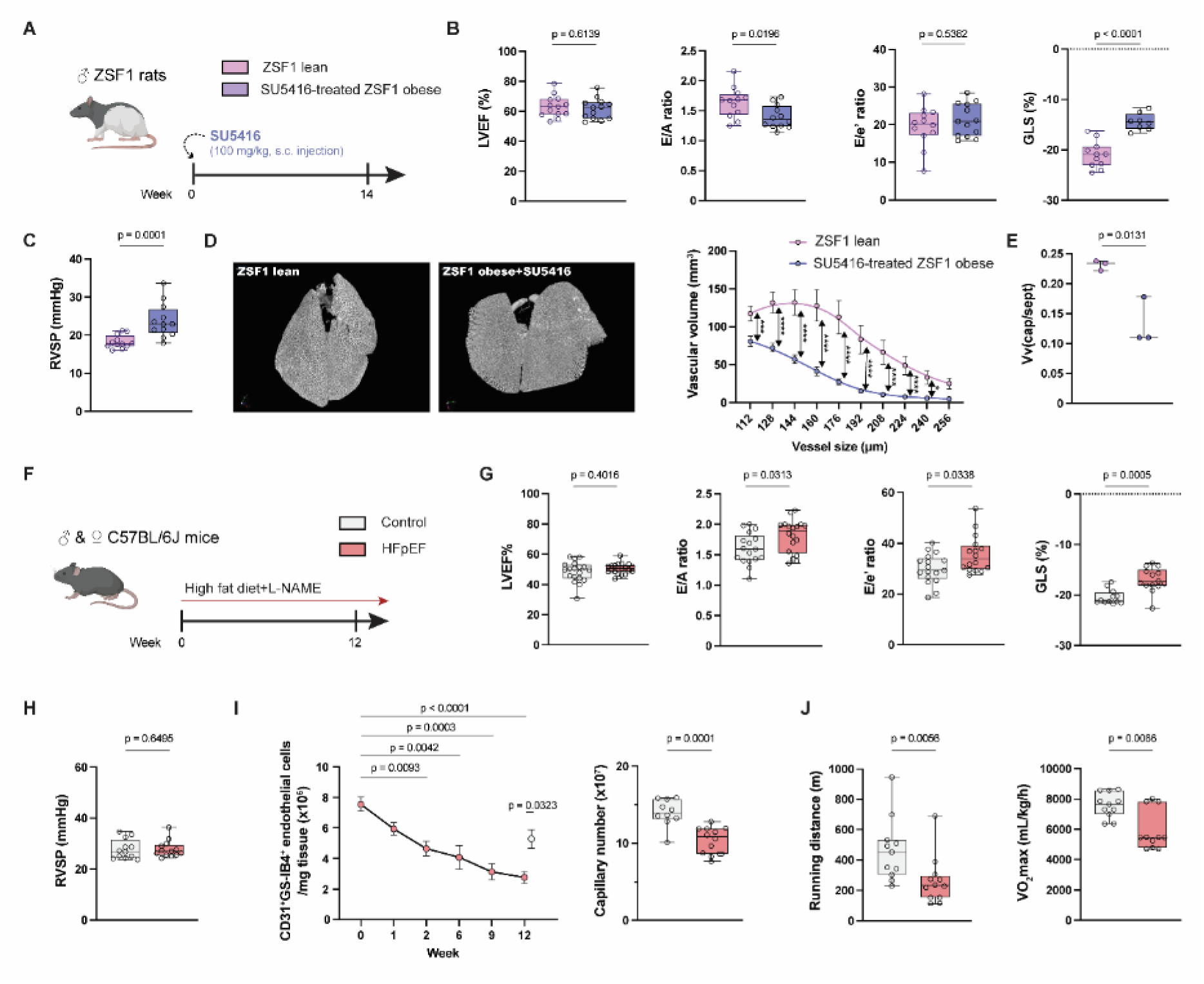
Lung microvascular rarefaction in HFpEF animals. **A**. Experimental protocol for HFpEF induction by SU5416 (s.c. 100 mg/kg) in ZSF1 obese rats. **B.** Echocardiographic analysis of left ventricular (LV) ejection fraction (LVEF), ratio of early to late LV filling velocity (E/A ratio), ratio of early diastolic mitral inflow velocity over early diastolic mitral annular tissue velocity (E/e’ ratio) and global longitudinal strain (GLS) in ZSF1 lean and SU5416-treated ZSF1 obese (n=13-15 each) rats. **C.** Right ventricular systolic pressure (RVSP) in ZSF1 lean and SU5416-treated ZSF1 obese rats (n=12 each). **D.** Representative micro-computed tomography (µCT) reconstructions of the pulmonary vasculature in ZSF1 lean (left) and SU5416-treated ZSF1 obese (right) rats and quantification of pulmonary vascular volume by µCT in vessels <250 µm in diameter in ZSF1 lean (n=4) and SU5416-treated ZSF1 obese (n=5) rats. Vessels <110 µm in diameter were excluded from the analysis due to limited resolution. **E.** Stereological quantification of capillary volume density within the interalveolar septa of the lung (V_v_(cap/sept)) in ZSF1 lean and SU5416-treated ZSF1 obese rats (n=3 each). **F.** Experimental protocol for HFpEF induction by high fat diet and Nω-Nitro-L-Arginin-Methylester-hydrochlorid (L-NAME) in *C57BL/6J* mice. **G.** Echocardiographic analysis of LVEF, E/A ratio, E/e’ ratio and GLS in control and HFpEF mice (n=15 each). **H.** RVSP in control and HFpEF mice (n=15 each). **I.** Flow cytometric quantification of CD31⁺/GS-IB4^+^ endothelial cells as a function of HFpEF progression over time (n=6 for each time point), and stereological quantification of total lung capillary number in control (n=10) and HFpEF (n=12) mice. **J.** Running distance and maximal oxygen consumption (VO₂max) of control and HFpEF mice (n=12 each) in treadmill exercise tests. Dots represent individual animals. Data are presented as median with interquartile range (IQR; all data except for D,I) or mean ± SEM (D,I). Mann–Whitney U-test (all data except for time course in I) or Kruskal-Wallis test followed by uncorrected Dunn’s multiple comparisons test; P-values as indicated or *p < 0.05; **p < 0.01; ***p < 0.001; ****p < 0.0001.

To further consolidate these results in a different species and to allow for subsequent mechanistic interrogation in genetically modified animals, we next probed for microvascular rarefaction in a murine HFpEF model of metabolic and hypertensive stress (Fig. 2F). In line with previous studies, the combination of high fat diet (HFD) with the nitric oxide synthase inhibitor L-NAME induced characteristic signs of diastolic dysfunction, reflected by a change in E/A ratio, an increased E/é ratio and a corresponding decrease in GLS (Fig. 2G). In line with a HFpEF phenotype, LVEF was again preserved (Fig. 2G), yet hemodynamic signs of overt PH were absent (Fig. 2H). Flow cytometric analyses revealed a time-dependent, exponential loss of lung microvascular endothelial cells and stereological assessment confirmed a reduced number of capillaries per lung in HFpEF mice (Fig. 2I). Finally, HFpEF mice showed characteristic signs of exercise intolerance, including reduced running distance and lower maximal oxygen consumption (VO₂max) during treadmill exercise (Fig. 2J). Full sets of echocardiographic, hemodynamic and gravimetric data as well as stereological data are given in Suppl. Tables 6&7.

### Lung microvascular rarefaction in AoB rats

To probe whether lung microvascular rarefaction is specific for HFpEF, we next characterized the pulmonary microvasculature in a preclinical heart failure model independent of prototypic triggers of HFpEF, namely AoB rats (Fig. 3A). After 9 weeks, AoB rats also showed signs of LV diastolic dysfunction including an increase in E/A and E/é ratio and a reduced GLS (Fig. 3B). While LVEF showed a slight trend to decline, this effect did not reach statistical significance. Similar to SU5416-treated ZSF1 obese rats, AoB rats had an increased RVSP (Fig. 3C). A full set of echocardiographic, hemodynamic and gravimetric data is given in Suppl. Table 8. µCT imaging did not detect significant differences in tissue or vascular volume at the total lung level between AoB rats and controls (Suppl. Fig. 2B); yet, lung vascular volume of vessels <250 µm in diameter was again significantly reduced consistent with loss of pulmonary microvessels (Fig. 3D). BD efflux analyses, flow cytometry (gating strategy shown in Suppl. Fig. 3) and quantitative stereology demonstrated a corresponding decrease in pulmonary capillary surface area, the total number of lung microvascular endothelial cells, and the total capillary number per lung (Fig. 3E), consolidating lung microvascular rarefaction in AoB rats. A full set of stereological parameters is given in Suppl. Table 9. Taken together, these findings identify pulmonary microvascular rarefaction as a characteristic pathological feature of left heart failure in different species and preclinical models.

**Figure 3.**
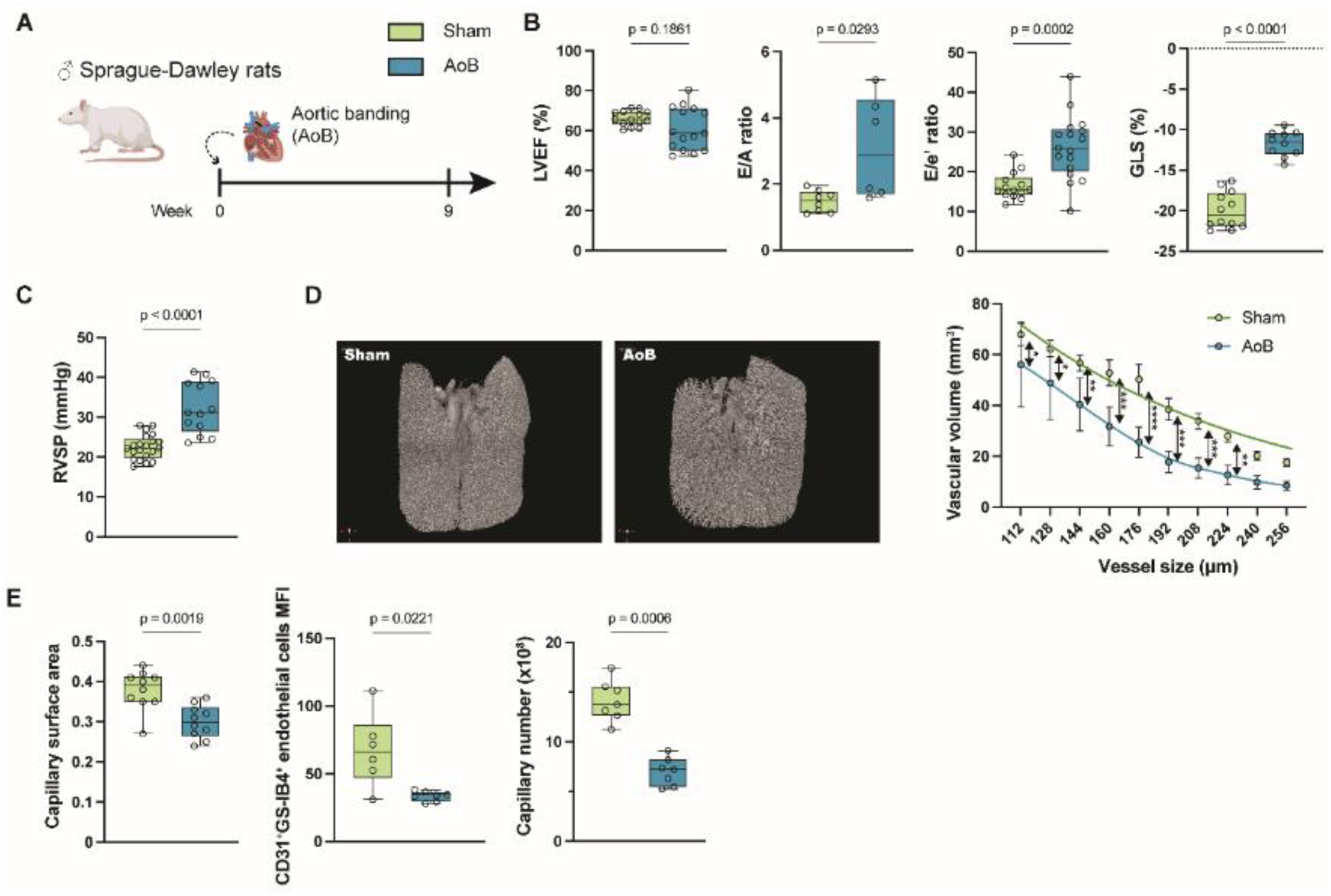
Lung microvascular rarefaction in AoB rats. **A**. Experimental protocol for left heart failure induction by aortic banding (AoB) in rats. **B.** Echocardiographic analysis of left ventricular (LV) ejection fraction (LVEF), ratio of early to late LV filling velocity (E/A ratio), ratio of early diastolic mitral inflow velocity over early diastolic mitral annular tissue velocity (E/e’ ratio) and global longitudinal strain (GLS) in sham (n=14) and AoB (n=16) rats. **C.** Right ventricular systolic pressure (RVSP) in sham (n=14) and AoB (n=16) rats. **D.** Representative micro-computed tomography (µCT) reconstructions of the pulmonary vasculature in sham (left) and AoB (right) rats and quantification of pulmonary vascular volume by µCT in vessels <250 µm in diameter in sham (n=4) and AoB (n=3) rats. Vessels <110 µm in diameter were excluded from the analysis due to limited resolution. **E.** Pulmonary capillary surface area in sham (n=10) and AoB (n=10) rats as determined by blue dextran efflux, flow cytometric quantification of CD31⁺/GS-IB4^+^ endothelial cells in sham (n=6) and AoB (n=7) rats, and stereological quantification of total lung capillary number in sham (n=7) and AoB (n=7) rats. Dots represent individual animals. Data are presented as median with interquartile range (IQR; all data except for D) or mean ± SEM (D). Mann–Whitney U-test; P-values as indicated or *p < 0.05; **p < 0.01; ***p < 0.001; ****p < 0.0001.

### HFpEF drives autophagy-dependent cell death in lung microvascular endothelial cells

In recent work, we identified chronic hypoxia to cause autophagy-driven cell death in lung microvascular endothelial cells^24^. To probe for a potentially similar scenario in HFpEF, we accessed a single cell RNA sequencing (scRNAseq) dataset from a previous study by Agrawal and coworkers^26^ which analyzed murine lungs in a similar model of HFD/L-NAME-induced HFpEF at an earlier timepoint (2 weeks). Unbiased clustering reproduced the same cell types as originally reported by the authors including different endothelial subsets, namely mesothelial, arteriolar, general capillary (gCap), aerocytes (aCap) and venular endothelial cells (Fig. 4A). Relative to controls, HFpEF mice showed a significant loss of gCaps while a concomitant decrease in aCaps did not reach significance (Fig. 4B). Conversely, the proportion of pericytes tended to increase. Consistently, an aggregate set of genes involved in regulated cell death pathways increased significantly in gCaps while aCaps did again not reach significance (Fig. 4C). This effect was associated with a parallel increase in an aggregate set of autophagy marker genes in both aCaps and gCaps, but not in pericytes (Fig. 4D). When compared at the single cell level, aggregate autophagy marker genes correlated with markers of regulated cell death pathways in gCaps (r_s_=0.38) and to a lesser extent also in aCaps (r_s_=0.26; Fig. 4E). These findings indicate a propensity for regulated lung endothelial cell death in HFpEF that is most pronounced in gCaps and less so in aCaps.

**Figure 4.**
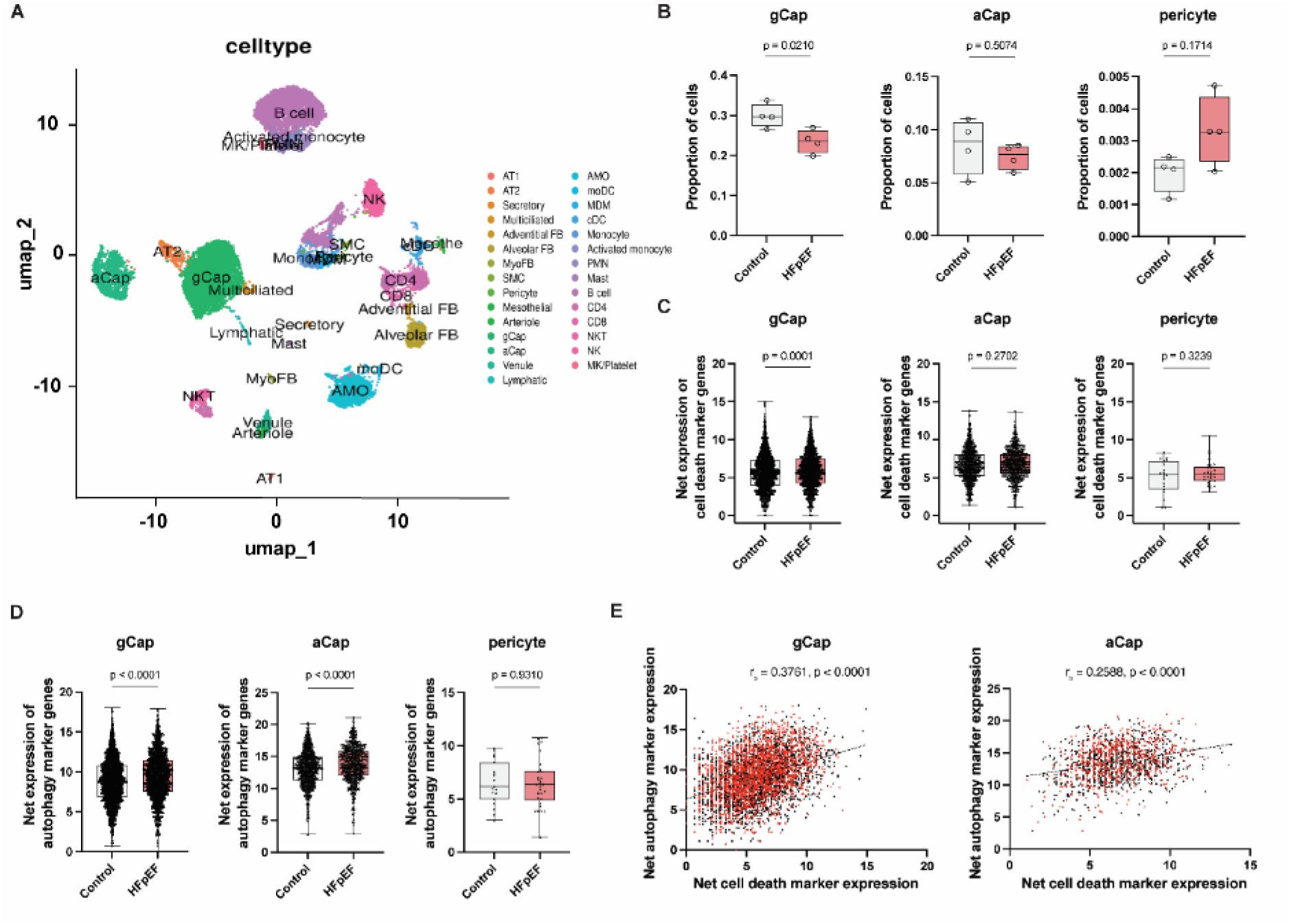
Single-cell RNA sequencing identifies propensity for regulated cell death and autophagy in general capillary cells. **A**. Uniform Manifold Approximation and Projection (UMAP) plot shows different cell clusters from integrated analysis of murine lungs after 2 weeks of HFpEF induction by high fat diet/L-NAME. **B.** Proportion of general capillary endothelial cells (gCap), aerocytes (aCap), and pericytes in HFpEF versus control mice (n=4 each). **C.** Net expression of an aggregate set of marker genes for regulated cell death pathway in gCap, aCap, and pericytes in HFpEF versus control mice (n=4 each). **D.** Net expression of an aggregate set of autophagy marker genes in gCap, aCap and pericytes in HFpEF versus control mice (n=4 each). **E.** Correlation analysis of autophagy marker genes vs. regulated cell death pathway marker genes in gCap and aCap. Dots represent individual animals (B) or cells (C-E), respectively. Data are presented as median with interquartile range (IQR). Mann–Whitney U-test; r_s_: Spearman coefficient of correlation; P-values as indicated.

Evidence for increased endothelial autophagy and regulated cell death was next validated at several levels in our HFpEF mouse model. First, we monitored autophagic flux in CD31^+^GS-IB4^+^ lung microvascular endothelial cells of HFpEF mice by flow cytometric detection of autophagic vacuoles with monodansylcadaverine. In line with our scRNAseq analyses, autophagic flux in lung endothelial cells increased within one week of HFpEF induction and reached its plateau between weeks 6 – 9 (Fig. 5A,B). Consistently, TEM identified endothelial autophagosomes in lung microvessels of HFpEF mice but not in control mice (Fig. 5C). Subsequent immunohistological analyses revealed an increased abundance of apoptotic endothelial cells, detected as CD31^+^ cells staining positive for cleaved caspase 3 (Fig. 5D) or cleaved-PARP1 (Fig. 5E), respectively, in HFpEF relative to control mice at week 12, and – in case of cleaved caspase 3 – as early as week 1.

**Figure 5.**
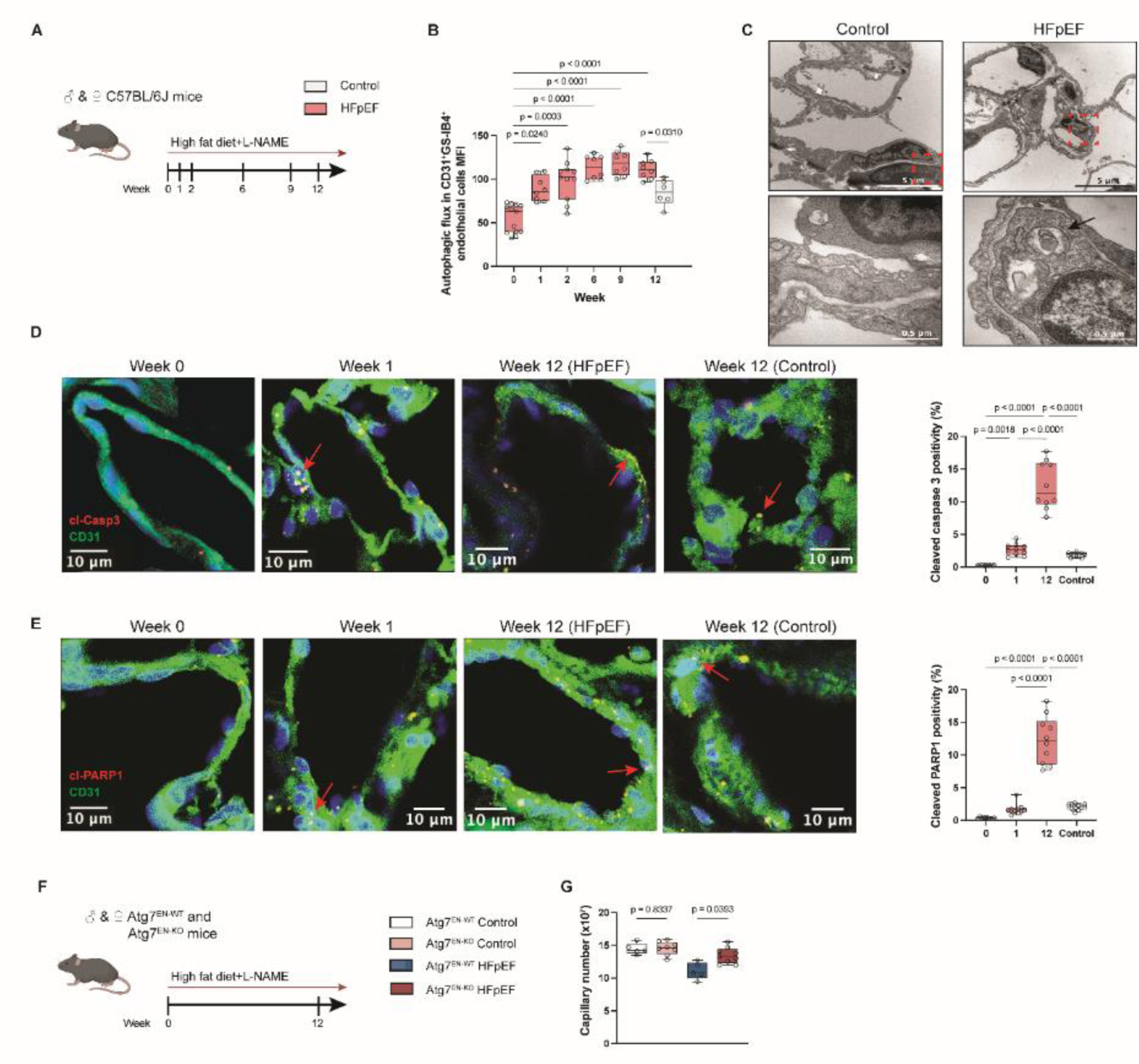
HFpEF drives autophagy-dependent cell death in lung microvascular endothelial cells. **A**. Experimental protocol for HFpEF induction by high fat diet and Nω-Nitro-L-Arginin-Methylester-hydrochlorid (L-NAME) in *C57BL/6J* mice. **B.** Flow cytometric analysis of autophagic flux in CD31⁺/GS-IB4^+^ lung microvascular endothelial cells as a function of HFpEF progression over time (n=6-13 for each time point). **C.** Representative transmission electron micrographs of lung endothelial cells at (from top to bottom) increasing levels of magnification showing autophagosomes (marked by black arrowheads) in HFpEF yet not in control mice. **D,E.** Representative images and quantitative analysis of cleaved caspase 3 (Cl-Casp3) (red; D) and cleaved poly[ADP-ribose] polymerase 1(Cl-PARP1) (red; E) staining in lung CD31^+^ endothelial cells (green; nuclei counterstained with DAPI in blue) from control (week 12) and HFpEF mice (week 0, 1, and 12) (n=10 each). Data are shown as percentage of vessels containing at least one Cl-Cas3 or Cl-PARP1 positive endothelial cell, respectively. **F.** Experimental protocol for HFpEF induction by high fat diet and L-NAME in endothelial cell-specific *Atg7*-deficient (*Atg7*^EN-KO^) and corresponding wild type (*Atg7*^EN-WT^) mice. **G.** Stereological quantification of total lung capillary number in *Atg7*^EN-WT^ (n=5 each) and *Atg7*^EN-KO^ (n=8 each) HFpEF or control mice. Dots represent individual animals. Data are presented as median with interquartile range (IQR). Kruskal-Wallis test followed by uncorrected Dunn’s multiple comparisons test; P-values as indicated.

To experimentally test for a functional role of increased endothelial autophagy in capillary rarefaction in HFpEF, we next subjected mice with an endothelial-specific knockout of the essential autophagy gene *Atg7* (*Atg7*^EN-KO^) to our murine HFpEF model (Fig. 5F). In stereological analyses, *Atg7*^EN-KO^ HFpEF mice had higher numbers of capillaries per lung relative to their corresponding *Atg7*^EN-WT^ controls, indicating that capillary rarefaction is prevented in the absence of excessive autophagic flux (Fig. 5G). A similar increase in capillary number in *Atg7*^EN-KO^ relative to *Atg7*^EN-WT^ mice was not evident under control conditions, consistent with the view that baseline autophagic flux does not cause excessive endothelial cell death.

### Systemic hypoxemia aggravates experimental HFpEF

To probe for functional effects of capillary rarefaction, we assessed parameters of gas exchange in both rat and mouse models. In AoB rats, microvascular rarefaction secondary to HF was associated with systemic hypoxemia as evidenced by reduced arterial partial pressure of oxygen (PaO_2_) and arterial oxygen saturation (SaO_2_), respectively (Fig. 6A,B). In contrast, venous partial pressure of oxygen (PvO_2_) remained unchanged (Fig. 6B), indicating impaired pulmonary gas exchange rather than increased peripheral oxygen extraction as cause of arterial hypoxemia. Analogous systemic hypoxemia was evident in HFpEF mice both at rest and after treadmill exercise (Fig. 6C,D). Conversely, attenuation of lung capillary rarefaction in *Atg7*^EN-KO^ mice with HFpEF improved oxygenation both at rest and after exercise (Fig. 6E,F), and this was associated with an extended running distance and improved maximal oxygen consumption (VO₂max; Fig. 6G) relative to *Atg7*^EN-WT^ mice. Notably, these effects were reversed in control, non-HFpEF mice, in that *Atg7*^EN-KO^ mice had worse oxygenation and exercise tolerance compared to their *Atg7*^EN-WT^ counterparts. As we will discuss below, these findings contrast the physiological relevance of baseline autophagy versus the pathological implications of excessive autophagy in HFpEF. Overall, these data identify pulmonary microvascular rarefaction as a consequence of excessive endothelial autophagy and cell death and as a characteristic pathological feature of heart failure that contributes to systemic hypoxemia and exercise intolerance.

**Figure 6.**
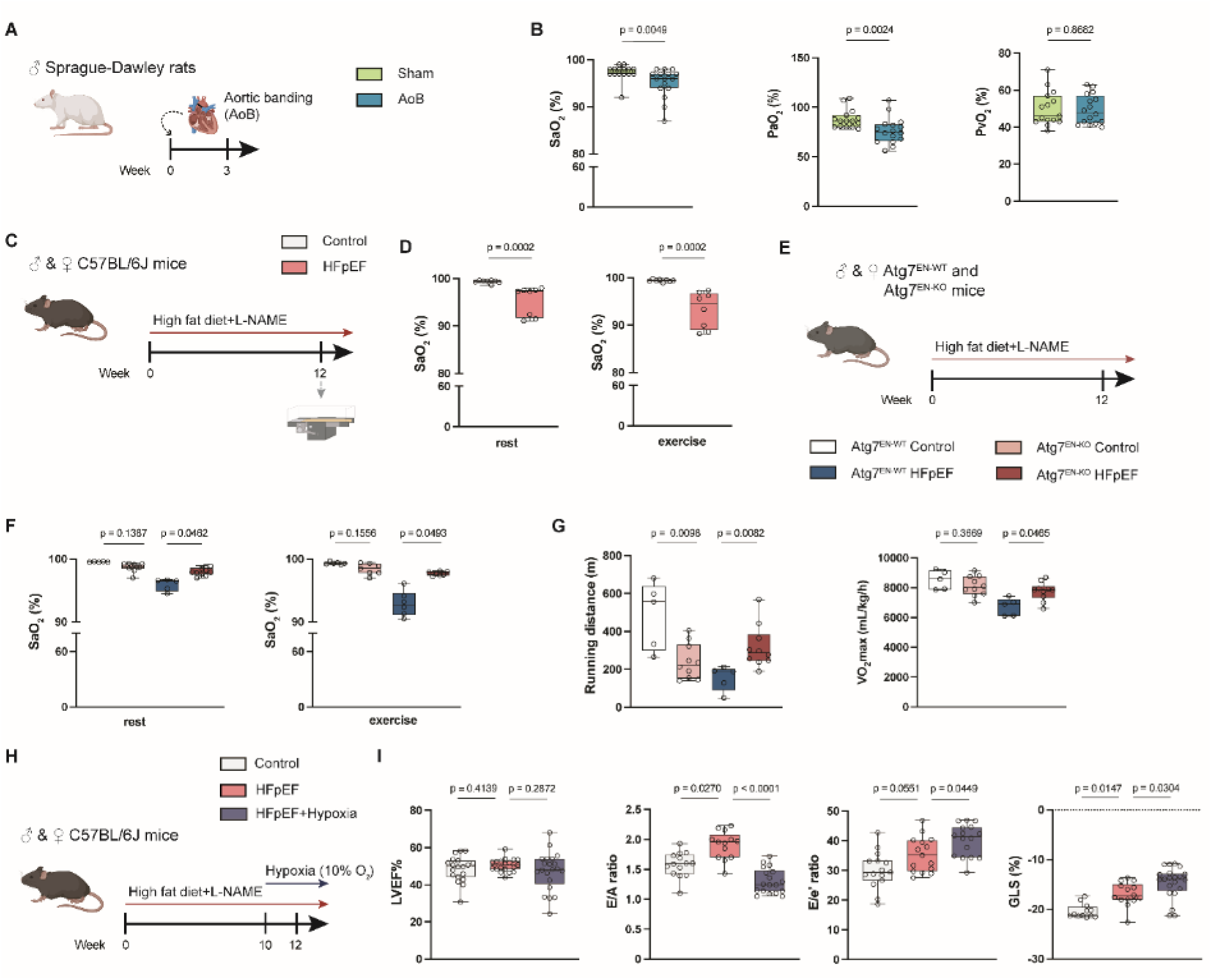
Systemic hypoxemia aggravates experimental HFpEF. **A**. Experimental protocol for induction of left heart failure by aortic banding (AoB) in rats. **B.** Arterial oxygen saturation (SaO₂), partial pressure of arterial oxygen (PaO₂), and mixed venous oxygen saturation (PvO₂) in sham (n=15) and AoB (n=16) rats. **C.** Experimental protocol for HFpEF induction by high fat diet and Nω-Nitro-L-Arginin-Methylester-hydrochlorid (L-NAME) in *C57BL/6J* mice. **D.** SaO₂ at rest and after exercise in control and HFpEF mice (n=8 each). **E.** Experimental protocol for HFpEF induction by high fat diet and L-NAME in endothelial cell-specific *Atg7*-deficient (*Atg7*^EN-KO^) and corresponding wild type (*Atg7*^EN-WT^) mice. **F.** SaO₂ at rest and after exercise in *Atg7*^EN-WT^ and *Atg7*^EN-KO^ mice (n=5-7 each). **G.** Running distance and maximal oxygen consumption (VO₂max) of *Atg7*^EN-WT^(n=5 each) and *Atg7*^EN-KO^ (n=10 each) HFpEF or control mice in treadmill exercise tests. **H.** Experimental protocol for hypoxia exposure (10%O_2_ for 2 weeks) in HFpEF mice. **I.** Echocardiographic analysis of left ventricular (LV) ejection fraction (LVEF), ratio of early to late LV filling velocity (E/A ratio), ratio of early diastolic mitral inflow velocity over early diastolic mitral annular tissue velocity (E/e’ ratio) and global longitudinal strain (GLS) in control, HFpEF and HFpEF+Hypoxia mice (n=15-21 each). Dots represent individual animals. Data are presented as median with interquartile range (IQR). Mann–Whitney U-test (B,D) or Kruskal-Wallis test followed by uncorrected Dunn’s multiple comparisons test (F,G,I); P-values as indicated.

To probe whether the resulting systemic hypoxemia may in turn affect severity and progression of heart failure, we next combined the HFpEF mouse model with a final 2-week exposure to hypoxia (10% O₂) starting after week 10 of HFD/L-NAME treatment (Fig. 6H). Echocardiographic assessment after 2 weeks of hypoxia revealed an exacerbated HFpEF phenotype with preserved LVEF but signs of aggravated LV dysfunction, indicated by an increased E/é and a further decrease in GLS while E/A ratio remained pathologically altered (Fig. 6I). Complete echocardiographic data are given in Suppl. Table 10. These findings demonstrate that hypoxemia can aggravate LV dysfunction, suggesting a potential detrimental feed-forward loop between HFpEF, lung capillary rarefaction and hypoxemia as driver of disease progression.

### Loss of endothelial autophagy and moderate hyperoxia mitigate experimental HFpEF

Contrasting the effects of hypoxia, prevention of capillary rarefaction and (partial) restoration of normoxemia in *Atg7*^EN-KO^ HFpEF mice was associated with improved LV diastolic function as demonstrated by lower E/A (yet without reaching significance at p=0.059) and E/é ratios as well as GLS relative to their corresponding *Atg7*^EN-KO^ controls (Fig. 7A,B). Complete echocardiographic data are given in Suppl. Table 11. To consolidate the link between restored normoxemia and improved LV function in a translationally relevant scenario, we finally tested whether moderate hyperoxia may similarly mitigate LV dysfunction in HFpEF mice. Analogous to our hypoxia experiments, we exposed HFpEF mice to 40% O₂ for the last 2 weeks of the HFD/L-NAME protocol (Fig. 7C). Similar to our findings in *Atg7*^EN-KO^ mice, hyperoxia normalized E/A and E/é ratios as well as GLS (Fig. 7D) while LVEF remained unchanged (Fig. 7D). Stereological assessment of lung tissue sections and arterial pulse oximetry at rest and after exercise further revealed that improved LV function was in turn associated with a restoration of lung capillarization and gas exchange (Fig. 7E,F). At the functional level, this resulted in restored exercise tolerance, evident as normalized running distance in treadmill tests (Fig. 7G). The complete set of echocardiographic data are given in Suppl. Table 12.

**Figure 7.**
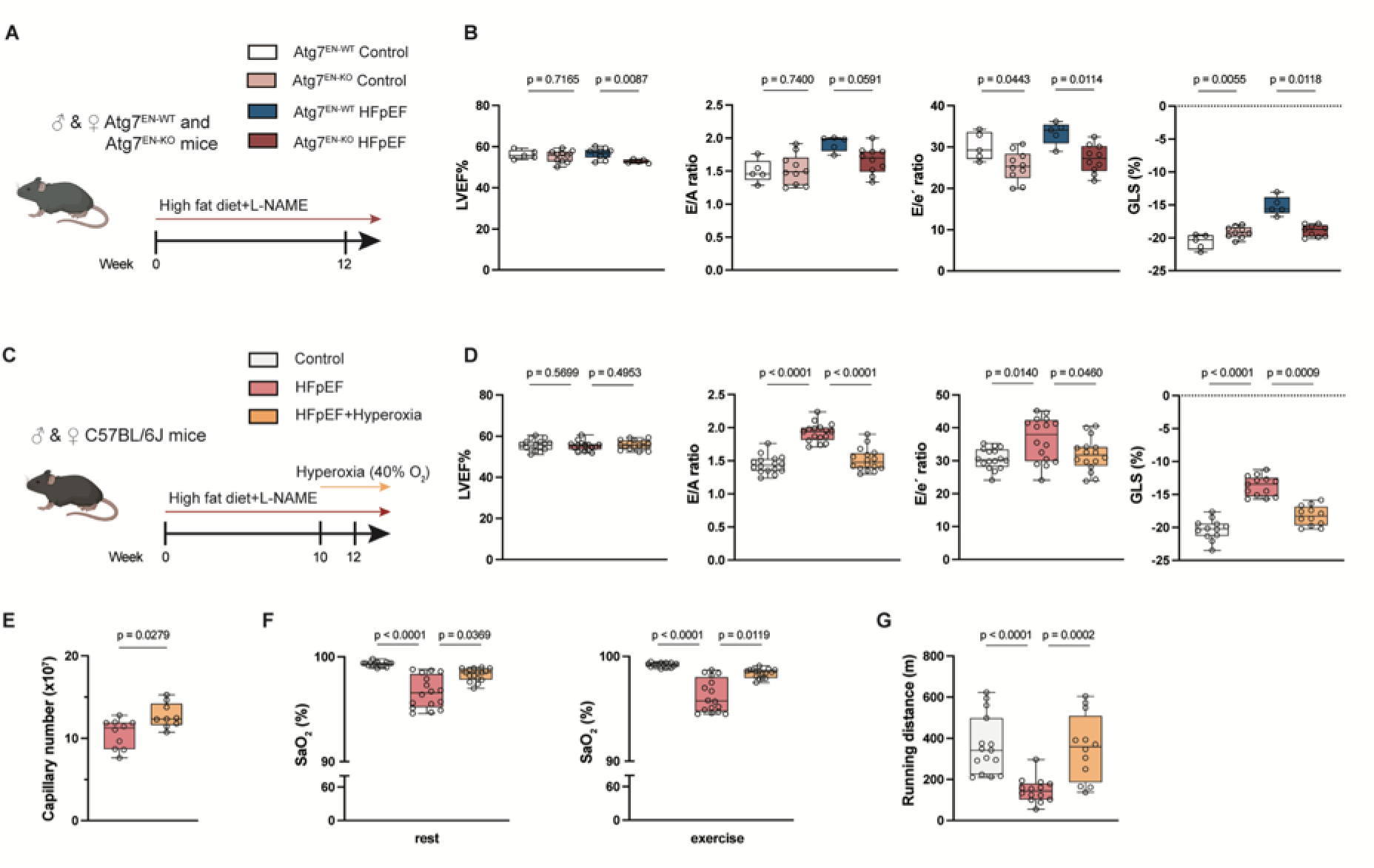
Loss of endothelial autophagy and moderate hyperoxia mitigate experimental HFpEF. **A**. Experimental protocol for HFpEF induction by high fat diet and Nω-Nitro-L-Arginin-Methylester-hydrochlorid (L-NAME) in endothelial cell-specific *Atg7*-deficient (*Atg7*^EN-KO^) and corresponding wild type (*Atg7*^EN-WT^) mice. **B.** Echocardiographic analysis of left ventricular (LV) ejection fraction (LVEF), ratio of early to late LV filling velocity (E/A ratio), ratio of early diastolic mitral inflow velocity over early diastolic mitral annular tissue velocity (E/e’ ratio) and global longitudinal strain (GLS) in *Atg7*^EN-WT^ (n=5 each) and *Atg7*^EN-KO^ (n=10 each) HFpEF or control mice. **C**. Experimental protocol for moderate hyperoxia exposure (40% O_2_ for 2 weeks) in HFpEF mice. **D.** Echocardiographic analysis of LVEF, E/A ratio, E/e’ ratio and GLS in control, HFpEF and HFpEF+Hyperoxia mice (n=16 each). **E.** Stereological quantification of total lung capillary number in HFpEF (n=10) and HFpEF+Hyperoxia (n=10) mice. **F.** Arterial oxygen saturation (SaO₂) at rest and after exercise in control, HFpEF and HFpEF+Hyperoxia mice (n=16 each). **G.** Running distance of control, HFpEF and HFpEF+Hyperoxia mice (n=16 each) in treadmill exercise tests. Dots represent individual animals. Data are presented as median with interquartile range (IQR). Mann–Whitney U-test (E) or Kruskal-Wallis test followed by uncorrected Dunn’s multiple comparisons test (all data except E); P-values as indicated.

## Discussion

In the present study, we identified in three different animal models of left heart disease extensive structural remodeling of pulmonary microvessels that was associated with impaired blood oxygenation and exercise performance. The latter findings were validated in a clinical HFpEF cohort in that disease severity and parameters of LV dysfunction correlated with impaired alveolo-capillary gas exchange. Mechanistically, microvascular rarefaction in HFpEF was driven by excessive autophagy and subsequent autophagy-associated cell death of lung capillary endothelial cells. At the functional level, the resulting hypoxemia aggravated LV dysfunction and thus, accelerated HFpEF progression. Conversely, moderate hyperoxia restored LV function, lung capillarization, gas exchange and exercise tolerance. These findings identify pulmonary microvascular rarefaction as a previously unrecognized pathological feature of HFpEF that drives disease progression and may contribute to recurrent oxygen desaturations and dyspnea in HFpEF patients (Fig. 8). Moderate oxygen therapy may present a promising avenue to decelerate or even reverse this disease trajectory.

**Figure 8.**
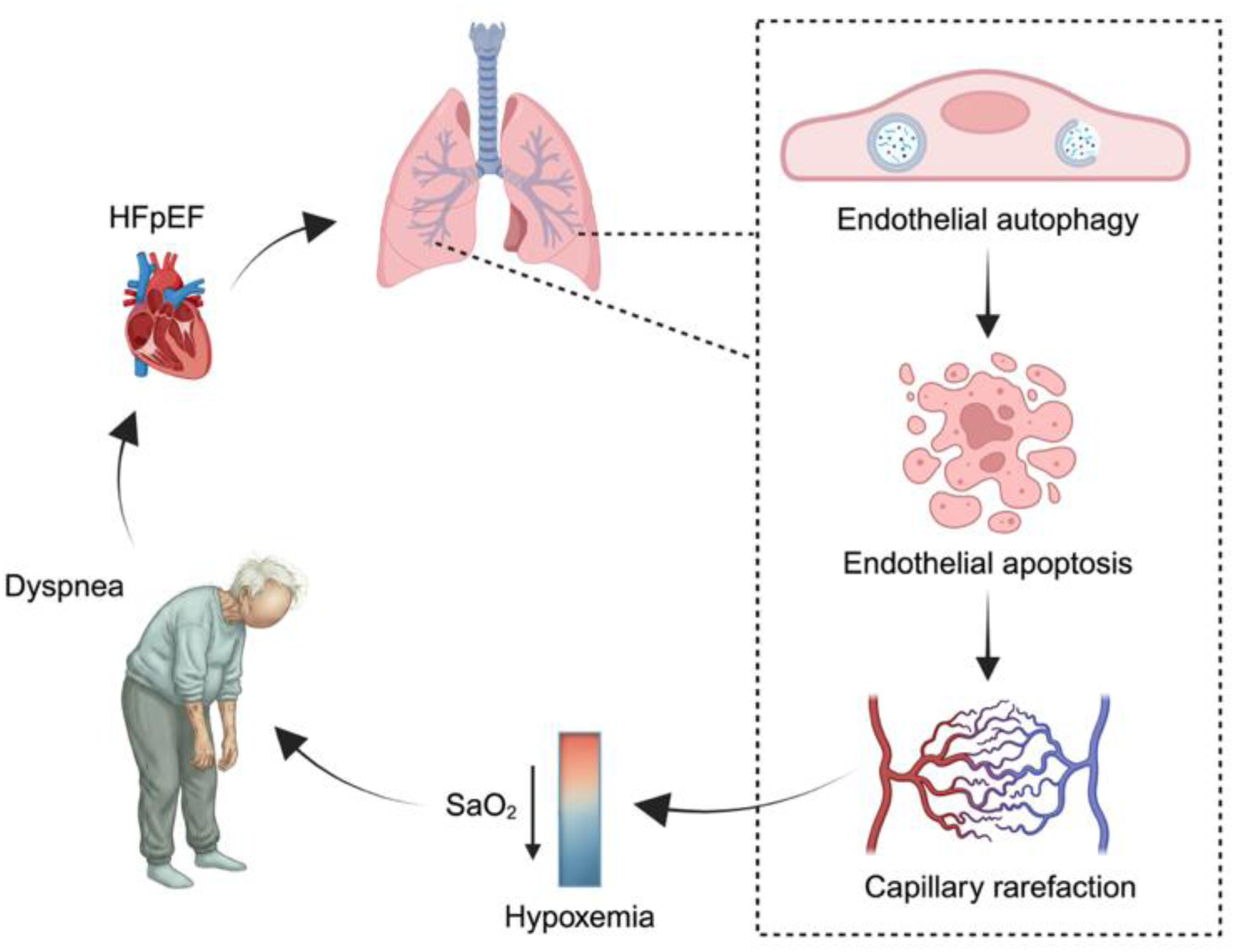
Graphical abstract. HFpEF causes lung microvascular rarefaction by driving autophagy-dependent endothelial cell death. Resulting hypoxemia contributes to clinical symptoms of dyspnea and aggravates HFpEF progression in a vicious circle.

### Impaired pulmonary gas exchange in HFpEF

Dyspnea or “air hunger” is one of the cardinal symptoms of HFpEF. Importantly, dyspnea is alternatively triggered via inputs from arterial chemosensors or lung mechanosensors^27^, rather than by changes in cardiac function *per se*. Indeed, arterial desaturations are frequently observed in HFpEF patients at rest and more characteristically during exercise^5,6^. This effect cannot be directly attributed to impaired hemodynamics in heart failure, as reduced CO and increased left atrial pressures do by themselves not decrease, but potentially even increase arterial oxygenation as they extend the contact time for blood with the oxygenation unit in the alveolo-capillary compartment^28^. Increased peripheral oxygen extraction due to low CO may potentially cause venous desaturation, which was, however, not evident in AoB rats and should be readily compensated for by the physiological oxygenation reserve capacity of the lung as long as alveolo-capillary gas exchange is uncompromised. As such, dyspnea and arterial hypoxemia in HFpEF point to structural alterations or functional deficiencies of the gas-exchange units in the lung.

Indeed, patients with HFpEF or other forms of heart failure frequently show characteristics of impaired alveolo-capillary gas exchange, evident as reduced DLCO^29–31^. Further expanding on these findings, we show for our clinical HFpEF cohort that arterial oxygen saturation at rest and during exercise, as well as DLCO decreased as a function of NYHA class, highlighting a direct association between disease severity and impaired pulmonary oxygen exchange. DLCO changes were independent of alveolar volume or hemoglobin concentration, consolidating the notion that impaired oxygenation and hypoxemia reflect a loss of effective alveolo-capillary diffusion capacity rather than changes in ventilated lung volume or blood oxygen-carrying capacity. Further, DLCO correlated with echocardiographic indices of diastolic dysfunction, indicating that declining efficiency of pulmonary gas exchange parallels progressive worsening of LV filling in HFpEF patients. As such, impaired pulmonary gas exchange emerges as a characteristic co-morbidity of HFpEF that not only parallels the severity of HFpEF, but – as we will discuss later – may also critically influence its progression.

Compromised lung diffusion capacity in HFpEF or other forms of heart failure has been alternatively attributed to the formation of cardiogenic edema^32^, structural thickening of the alveolo-capillary barrier^33^, and/or ventilation-perfusion mismatching^9,34–36^. The extent of both interstitial and alveolar edema in chronic heart failure patients is, however, relatively modest due to a series of effective adaptive, counter-regulatory mechanisms such as a reduction in endothelial permeability and an increase in alveolar fluid clearance^37,38^. Similarly, definite data showing remodeling of the alveolo-capillary barrier or impaired ventilation-perfusion matching in HFpEF patients and their contribution to impaired pulmonary gas exchange are so far lacking.

In the present work, we provide first evidence for a decline in lung vascular gas exchange surface area as pathomechanism for impaired alveolo-capillary gas exchange in HFpEF. While surface area as a key determinant of alveolo-capillary gas diffusion is commonly discussed in terms of alveolar surface, it is important to consider the equal relevance of capillary surface area as its vascular counterpart. In the present study, lung microvascular rarefaction was evidenced in three different animal models of heart failure in two different species –SU5416-treated ZSF1 obese rats, HFD/L-NAME-treated mice, and AoB rats – by a set of different methodologies: µCT revealed a loss of lung vascular volume in rat lungs that was most pronounced in the microvascular compartment comprised of vessels <250 µm in diameter. As µCT can, however, not resolve single alveolar capillaries, we consolidated microvascular rarefaction at the capillary level by quantitative stereology, commonly considered as the gold standard of lung morphometry^39^. Specifically, we calculated the total number of capillaries per lung from the Euler-Poincaré characteristic based on recognizable topological constellations termed “islands”, “bridges” and “holes” in an unbiased manner^20^. Compared to their respective controls, both HFpEF mice and AoB rats showed a significant reduction in total capillary number per lung, consolidating that microvascular rarefaction extended to the functional units of gas exchange. Consistent with capillary rarefaction, flow cytometric analyses revealed a time-dependent loss of microvascular EC in lungs with the progression of HFpEF in mice that exceeded 50% after 12 weeks of HFD and L-NAME treatment. Microvascular EC loss of similar magnitude was evident in AoB rats after 9 weeks. In AoB rats, we further consolidated the actual loss of capillary gas exchange area by measuring the efflux over time of a dye previously loaded to capillary EC in isolated perfused lungs^25^. As such, we provide multiple levels of evidence for a structural loss of functional gas exchange area in different animal models. The fact that microvascular rarefaction was equally evident in classic rodent models of HFpEF as well as in a metabolic stress-independent model of pressure overload in AoB rats highlights loss of pulmonary gas exchange units as universal feature of heart failure, suggesting in turn that microvascular rarefaction is primarily driven by changes in pulmonary hemodynamics common to different forms of heart failure rather than by their individual pathogenetic mechanisms. In line with this notion, clinical finding of reduced DLCO and symptoms of dyspnea span across the entire spectrum of heart failure patients^29–31^.

The extent of EC loss required to impair alveolo-capillary gas exchange is presently unknown. The network of alveolar capillaries is extremely dense; that notwithstanding any decrease in capillary gas exchange area will directly translate into a proportional decrease in oxygen diffusion as described by Fick’s first law of diffusion. At rest, impaired diffusion may in part be compensated by the fact that blood oxygenation is completed before the blood exits the pulmonary capillary bed^40^, yet these compensatory mechanisms tend to fail with shorter capillary transit times as seen e.g. in exercise. In line with this concept, arterial desaturation in patients with NYHA class II or III became more prominent during exercise as compared to rest.

(Micro)vascular rarefaction has previously been reported as a characteristic feature in patients with and animal models of pulmonary arterial hypertension^24,41,42^. To our knowledge, the present work constitutes, however, the first report of a similar phenomenon in left heart disease. This finding is unexpected as the increased left atrial pressures seen in HF will acutely recruit previously unperfused alveolar capillaries and further distend open microvascular segments^28^, resulting in a net increase in capillary gas exchange surface. Our present data show, however, that this acute effect is subsequently counteracted by a progressive loss of EC and lung capillaries.

Further mechanistic interrogation revealed increased autophagic flux and a higher propensity for apoptotic cell death in lung microvascular endothelial cells of HFpEF mice as compared to controls. Specifically, analysis of a single-cell transcriptomic dataset of murine lungs at an early stage of HFpEF (2 weeks after start of HFD/L-NAME) yielded increased expression of regulated cell death pathway and autophagy related genes predominantly in gCaps that was associated with a loss in total gCap cell numbers. The fact that these effects were more prominent in gCaps than aCaps may at first seem counterintuitive in the context of HFpEF associated arterial desaturation, as aCaps form the primary vascular unit for pulmonary gas exchange^43^. However, as gCaps serve as progenitor pool for the largely non-proliferative aCap population^44^, loss of gCaps at an early stage (shown in the present scRNAseq dataset to occur within 2 weeks of HFpEF) is expected to have profound effects also on aCaps over the subsequent progression of HFpEF. Consistently, flow cytometric and immunohistological analyses of lungs from HFpEF mice revealed a progressive increase in autophagic flux over time and elevated markers of endothelial apoptosis, specifically cleaved caspase-3 and cleaved-PARP1 in CD31^+^ cells at 12 weeks HFD/L-NAME. While physiological autophagy constitutes an important cell survival mechanism, an emerging body of evidence shows that excessive autophagy can trigger apoptotic cell death (autophagy-dependent cell death, ADCD)^45^. In the present study, this dichotomy likely accounts for the finding that control mice with EC-specific loss of the essential autophagy gene *Atg7* only managed a shorter running distance compared to their wild type counterparts in treadmill tests, yes performed better in the HFpEF model.

In previous work, we reported ADCD as a driver of lung microvascular EC loss in chronic hypoxic PH^24^. In the present study, we identified an analogous mechanism in HFpEF, in that deficiency of the essential autophagy gene *Atg7* protected HFpEF mice from lung EC apoptosis and capillary loss. Notably, this finding is in line with the results from a cardiome-directed network analysis of RNAseq data in HFpEF rats which identified apoptosis and autophagy as one of five core disease processes^46^. As such, excessive autophagy and ADCD may constitute key drivers of both heart and lung disease in HFpEF. In parallel, ADCD of lung microvascular endothelial cells emerges as a principal characteristic across different types of PH (namely PH due to chronic hypoxia or HFpEF).

Importantly, the development of lung capillary rarefaction and subsequent hypoxemia promote the progression of ventricular dysfunction in HFpEF in a detrimental feedforward loop. Exposing HFpEF mice to hypoxia worsened characteristic signs of LV diastolic dysfunction, while moderate hyperoxia improved LV function and exercise tolerance. Notably, hyperoxia also restored pulmonary capillarization and normalized arterial oxygenation, indicating that the “vicious circle” between HFpEF and lung capillary rarefaction can be reversed into a “virtuous circle”. As such, ambulatory oxygen therapy e.g. by high flow nasal oxygen may not only alleviate symptoms of dyspnea and increase exercise capacity in HFpEF patients^47^, but may provide a therapeutic means to decelerate disease progression. Beyond oxygen therapy, interventions targeting the underlying mechanisms of microvascular rarefaction may guide future therapeutic strategies with the aim to reduce endothelial apoptosis and/or modulate excessive endothelial autophagy to preserve lung microvascular integrity.

Several methodological aspects and limitations of the present study merit specific consideration. First, pressure-overload models are commonly considered as HFrEF rather than HFpEF models. Hence, although the majority of our AoB rats had an EF >50% and showed signs of diastolic dysfunction such as increased E/A, E/é and a reduced GLS, caution is required in the interpretation of this model as HFpEF, as it may transition to a HFrEF phenotype at later stages. In the present manuscript, we thus refer to AoB rats as a model of heart failure due to pressure overload. In contrast, the SU5416-treated ZSF1 obese rat and the HFD/L-NAME murine model present well-established models of LV HFpEF, which was confirmed in the present study by echocardiographic signs of diastolic dysfunction in the presence of a sustained EF. Importantly, microvascular rarefaction and impaired gas exchange were consistent across models irrespective of LVEF, suggesting that this pulmonary phenotype reflects fundamental consequences of chronic pulmonary congestion secondary to LV failure rather than a response unique to a specific subtype of HF. Second, E/A ratios tended to change in different directions across models, with E/A increasing in HFpEF patients, AoB rats, and HFD/L-NAME mice while they decreased in SU5416-treated ZSF1 obese rats and in hypoxia-exposed HFD/L-NAME mice. This seemingly discrepant effect is well-recognized in the literature: As the transmitral flow pattern depends both on loading conditions and diastolic function, E/A may decrease in HF due to impaired relaxation or increase due to restrictive filling. Importantly, all models exhibited clear evidence of diastolic dysfunction based on changes in GLS and/or E/e′, validating E/A as an informative but context-dependent indicator of diastolic abnormalities. Third, in ZSF1 rats image quality and the resolution at light microscopic level was not sufficient to allow for the stereological assessment of capillary number per lung. We thus adopted an alternative, TEM based stereological approach to estimate capillary volume density within the interalveolar septa of the lung in these animals. The identified loss of capillary structures was nevertheless internally consistent in both quality and quantity with results obtained by classical stereology in AoB rats or HFpEF mice. Fourth, it should be emphasized that the degree of hypoxia used (10% O₂) exceeds the level of hypoxemia typically observed in human HFpEF. The results of this intervention should therefore be interpreted primarily as proof-of-principle demonstrating the impact of impaired gas exchange on cardiac function rather than as a direct clinical analogue. Finally, it should be noted that in HFpEF mice, capillary rarefaction was not associated with a corresponding increase in RVSP. This observation aligns with prior work by Duncan Stewart and colleagues showing that RVSP only starts to increase after more than 2/3 of the pulmonary microvascular bed has been lost^48,49^. As such, impairments in gas exchange may precede overt PH in the trajectory of HFpEF.

In summary, our clinical and experimental data converge on a new and previously unrecognized mechanism of exertional dyspnea in heart failure with important functional and therapeutic consequences for disease progression. Through autophagy– and apoptosis-driven loss of lung endothelial cells, HFpEF causes a significant loss in surface area for alveolo-capillary gas exchange, thus contributing to impaired diffusion capacity, systemic hypoxemia and exercise intolerance. Hypoxemia, in turn, aggravates LV dysfunction, thereby establishing a feed-forward loop between cardiac and pulmonary pathology. The ability of moderate hyperoxia or inhibition of endothelial autophagy to restore not only blood oxygenation, but also pulmonary capillarization and – importantly – improve cardiac diastolic performance highlights the therapeutic potential of targeting the lung microvasculature and/or systemic oxygenation in heart failure. These findings broaden the mechanistic framework of heart failure beyond the myocardium and identify the pulmonary microcirculation as a critical — and potentially modifiable — determinant of disease progression.

## Acknowledgement

The authors thank all Kuebler and Grune lab members for discussion and insightful comments. We thank Kerstin Riskowski, John Horn, and Katja Dörfel for technical assistance and Dr. Sara Timm and Petra Schrade from the Core Facility of Electron Microscopy for support. We thank the Advanced Medical Bioimaging Core Facility (AMBIO) of the Charité for support in acquisition of imaging data. The graphical abstract has been created in BioRender. Kocana, C. (2026) https://BioRender.com/rio5nw2. Dyspnea figure in the graphical abstract was created with Illustrae.co.

## Author contribution

C.K. and L.J. performed and analyzed experiments and interpreted data. C.K. made the figures. N.H., P.S., Q.L., K.K., A.M., D.F., L.v.O., P.-L.P., J.L.G. and A.W. performed experiments and collected data. A.M.C., K.F., E.R., A.K., L.K., D.Z. and V.Z. provided human specimen and data. S.M.T., M.A.H., B.K.B. and S.A.H. performed single-cell RNA sequencing analysis. J.G. and W.M.K. designed experiments. M.U., C.B., J.G. and W.M.K. discussed results and strategy. C.K. and W.M.K. wrote the manuscript with input from all authors. W.M.K. conceived and directed the study.

